# Evolutionary tracking of SARS-CoV-2 genetic variants highlights an intricate balance of stabilizing and destabilizing mutations

**DOI:** 10.1101/2020.12.22.423920

**Authors:** Jobin John Jacob, Karthick Vasudevan, Agila Kumari Pragasam, Karthik Gunasekaran, Balaji Veeraraghavan, Ankur Mutreja

## Abstract

The currently ongoing COVID-19 pandemic caused by SARS-CoV-2 has accounted for millions of infections and deaths across the globe. Genome sequences of SARS-CoV-2 are being published daily in public databases and the availability of this genome datasets has allowed unprecedented access into the mutational patterns of SARS-CoV-2 evolution. We made use of the same genomic information for conducting phylogenetic analysis and identifying lineage-specific mutations. The catalogued lineage defining mutations were analysed for their stabilizing or destabilizing impact on viral proteins. We recorded persistence of D614G, S477N, A222V V1176F variants and a global expansion of the PANGOLIN variant B.1. In addition, a retention of Q57H (B.1.X), R203K/G204R (B.1.1.X), T85I (B.1.2-B.1.3), G15S+T428I (C.X) and I120F (D.X) variations was observed. Overall, we recorded a striking balance between stabilizing and destabilizing mutations, therefore well-maintained protein structures. With selection pressures in the form of newly developed vaccines and therapeutics to mount soon in coming months, the task of mapping of viral mutations and recording of their impact on key viral proteins would be crucial to pre-emptively catch any escape mechanism that SARS-CoV-2 may evolve for.

**STUDY IMPORTANCE:** As large numbers of the SARS CoV-2 genome sequences are shared in publicly accessible repositories, it enables scientists a detailed evolutionary analysis since its initial isolation in Wuhan, China. We investigated the evolutionarily associated mutational diversity overlaid on the major phylogenetic lineages circulating globally, using 513 representative genomes. We detailed phylogenetic persistence of key variants facilitating global expansion of the PANGOLIN variant B.1, including the recent, fast expanding, B.1.1.7 lineage. The stabilizing or destabilizing impact of the catalogued lineage defining mutations on viral proteins indicates their possible involvement in balancing the protein function and structure. A clear understanding of this mutational profile is of high clinical significance to catch any vaccine escape mechanism, as the same proteins make crucial components of vaccines recently approved and in development. In this direction, our study provides an imperative framework and baseline data upon which further analysis could be built as newer variants of SARS-CoV-2 continue to appear.

## INTRODUCTION

The emergence of severe acute respiratory syndrome coronavirus 2 (SARS-CoV-2) in Wuhan, China and the subsequent global spread has brought the world to a standstill (1). During the course of 11 months the coronavirus disease 19 (COVID-19) pandemic has caused more than 81 million confirmed cases in 220 countries with close to 1,770,000 fatalities. (2). Initially, and rightly too, the efforts were focused on minimising the number of cases and deaths due to COVID-19 (3). This included fast tracking the search and development of novel treatment and prevention options (4). Today, however, as vaccine candidates have started showing promising results, there is a cautious shift towards assessing the efficacy of vaccine candidates with respect to the circulating diversity of SARS-CoV-2 and its continuously evolving genetic variants (5).

Functional mutations that help the virus to adapt to the recent host-shift events are hypothesised to drive the evolution of transmissibility and virulence in SARS-CoV-2 (6). Shortly after the first isolated SARS-CoV-2 genome from China was published, >30,500 distinct mutations were catalogued in the CoV-GLUE (http://cov-glue.cvr.gla.ac.uk/) among globally circulating strains of this virus (7). Variation in the genetic makeup are key determinants in measuring the evolutionary distance and stability of SARS-CoV-2 from the first sequenced isolate (8). Moreover, tracking the evolution of SARS-CoV-2 since its introduction in humans is a high priority undertaking to prevent future waves of this pandemic from escaping the global preparedness (9). Since many vaccine candidates currently under development are derived from the first available SARS-CoV-2 sequences, recurrent genetic changes may have an unforeseen impact on their sustained effectiveness in the longer term (10).

The availability of whole genome sequences of SARS-CoV-2 in public repositories such as Global Initiative on Sharing All Influenza Data (GISAID) and real-time data visualisation pipeline NextStrain (https://nextstrain.org) offers a great opportunity for scientists to track the evolutionary path of this virus (11, 12). Phylogenetic Assignment of Named Global Outbreak LINeages tool (PANGOLIN) has been the most widely used tool for lineage assignment to newly emerging variants. PANGOLIN (https://cov-lineages.org/pangolin.html) has also been deployed in establishing the transmission patterns of various clones of this virus (13). Since coronaviruses frequently recombine, tracking the evolution and assigning lineages has been challenging (13, 14). As a result, multiple studies that tracked the evolution of SARS-CoV-2, have been hugely controversial. For example, doubts have been cast on the claim of finding more aggressive L type emerging from S type strains (14). Similarly, the hypothesis of rapid spread of D614G variant of SARS-CoV-2 indicating a possible fitness advantage has been questioned (15 - 17). Therefore, in the current and highly sensitive global circumstances due to this pandemic, having a detailed map of mutations highlighting their prospective role in therapeutics and vaccine development can prepare us better for the future waves of continuously evolving SARS-CoV-2. In this study, we present a catalogue of the most important genomic mutations recorded between December 2019 – November 2020 in SARS-Cov-2 and their possible impact on stability of protein candidates that form crucial part of vaccines and also constitute the most common therapeutic targets.

## MATERIAL AND METHODS

### Data acquisition and curation

In total, we have retrieved 7000 genomes from GISAID EpiCoV database (https://www.gisaid.org/). Datasets that were flagged as complete (>28,000 bp) were screened and subsequently manually curated for excluding low quality/coverage sequences and duplicates. Sequence metadata was retrieved and genome containing sampling time and location were only chosen for the study. Lineages were assigned from alignment file using the Phylogenetic Assignment of Named Global Outbreak LINeages tool PANGOLIN v1.07 (https://github.com/hCoV-2019/pangolin). We selected a subset of 513 genomes (Supplementary Table S1) that belongs to all major PANGOLIN lineages and common mutations for the optimal output of the phylogenetic tree.

### Phylogenetic analysis

Genome sequences were aligned against the original Wuhan-Hu-1 genome (Accession: NC_045512) using multiple genome sequence alignment tool MAFFT (v6.240) (18). Subsequently, the error prone 5’-UTR and 3’-UTR regions were masked and the genome size was adjusted without losing key sites. Maximum likelihood tree was generated using IQTREE v.1.6.1 (http://www.iqtree.org/) under the GTR nucleotide substitution model with 1000 bootstrap replicates (19). The ML tree was visualised and labelled using the interactive tree of life software iTOL v.3 (20).

### Mutation Profiling

In order to identify the genetic variants, assembled genomes were mapped against the reference (Wuhan-Hu-1: Accession: NC_045512) using Snippy mapping and variant calling pipeline (https://github.com/tseemann/snippy) (21). Among the SNPs, missense SNPs (nonsynonymous) was extracted using custom written bash scripts and manually curated as per CoV-GLUE database (http://cov-glue.cvr.gla.ac.uk/). Specifically, we considered eleven lineage defining mutations and 59 major missense mutations in four major structural proteins: Envelope protein (E), Membrane glycoprotein (M), Nucleocapsid phosphoprotein (N) and Spike protein (S). Structural analysis of these 70 amino acid substitutions in SARS CoV-2 mutants were analysed to examine the potential impact of these mutations on protein stability.

### Structural analysis

The structural impact of mutations has been assessed from COVID-3D server (http://biosig.unimelb.edu.au/covid3d) which has integrated analytics regarding mutation-based structural changes in a protein. Vibrational entropy/VE (ΔΔS) and unfolding Gibbs free energy/FE (ΔΔG) were considered as markers to ascertain the stability of the variants. Gibbs free energy/FE (ΔΔG) values from Site Directed Mutator (SDM), DUET and DynaMut tools available in COVID-3D server were considered (22, 23). The change in vibrational entropy energy (ΔΔSVib ENCoM) between wild-type and mutant protein was calculated using DynaMut (24). VE explains the occupation probabilities of protein residues in an energy landscape based on average configurational entropies. Considerable decrease in VE increases the rigidity of the proteins (25). FE on the other hand describes the free energy alterations while unfolding a kinetically stable protein (24). The positive and negative values of ΔΔG indicate the stabilizing and destabilizing mutations. DynaMine (http://dynamine.ibsquare.be/) was employed to validate the stability profiles through residue level (sequence-based) dynamics. Backbone N-H S2 order parameter values (atomic bond vector’s movement restrictions) were generated according to the molecular reference frame. These N-H S^2^ order parameter values are evaluated from experimentally determined NMR chemical shifts. A value above 0.8 is considered as highly stable, values between 0.6-0.8 can be considered to be functionally contextual, and values >0.6 are highly flexible (26).

## RESULTS

### Diversity of SARS-CoV-2 Genomes

Of the 7000 SARS-CoV-2 genomes screened, we constructed a robust phylogenetic tree on strategically selected 513 genomes that reflected the most complete diversity among the isolates by covering all the PANGOLIN lineages. Lineage assignment based on PANGOLIN tool indicated the circulation of seven distinct lineages and/or sub-lineages such as A, B.1, B.1.1, B.1.1.1, B.2, B.3, B.4 and B.6. This is in line with the phylogenetic groupings by GISAID (S, L, V, O, G, GH and GR) (Figure 1). As the epidemic has progressed and mutations have accumulated, further subdivision of major lineages into sub-lineages has been observed. Overall, a total of 61 lineages and sub-lineages have been found to be circulating concurrently in multiple countries around the world. In general, numerous introductions of different variants were observed across the globe with a few sub-lineages (C.2, D.2) being restricted to certain regions. While B.1.113 lineage, for example, has been exclusively reported from India, lineages C.2 and D.2 geographically have been confined to South Africa and Australia, respectively.

**Figure 1.**
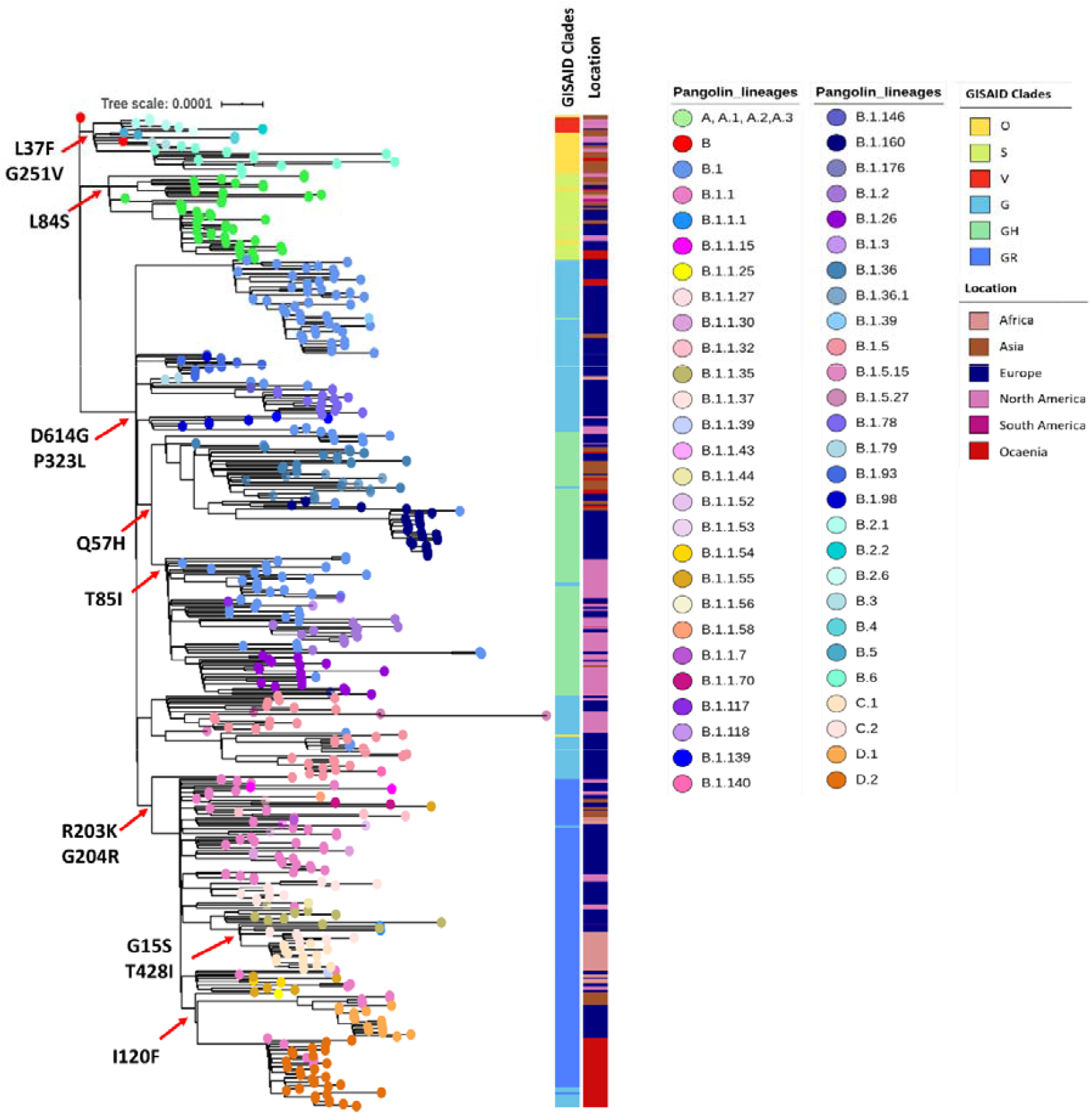
Maximum Likelihood phylogenetic tree inferred from 513 SARS-CoV-2 genomes. The tree was constructed using multiple genome sequence alignment (MAFFT) by mapping against the Wuhan-Hu-1 strain (Accession: NC_045512). Tips are coloured the major lineages assigned by PANGOLIN. Respective lineages assigned by GISAID and origin of sequence are labelled as colour strips. The scale bar indicates the distance corresponding to substitution per site.

### Major amino acid substitutions

Mutation mapping showed a total of 106 amino acid substitutions (missense mutations in >5 genomes) from a representative set of 513 genomes. The analysis also revealed 36 mutations that were found in >5% of genome sequences while 12 major substitutions were lineage defining mutations (Figure 1). The first major mutation to appear was L84S in ORF8, (present in 8.6% of the genomes) that has defined A lineage (i.e., clade S in GISAID classification). The subsequent amino acid substitutions L37F in ORF3a and G251V in nsp6 were found to be present in 13.3% and 1.4% of genomes, respectively. The combination of G251V and L37F, which was initially considered as a defining mutation pattern for B.2 – B.6 lineage (clade V in GISAID classification), more detailed analysis has shown that isolates carrying G251V mutation are distributed in other lineages too. The predominant lineage defining mutations in the whole dataset were D614G (85.5%) and P323L (85.5%), after originally appearing in late January 2020 (Figure 2). Other major mutations noted are Q57H (26.5%), R203K/G204R (33%), G15S (12%), I120F (11.5%) and T85I (14%).

**Figure 2.**
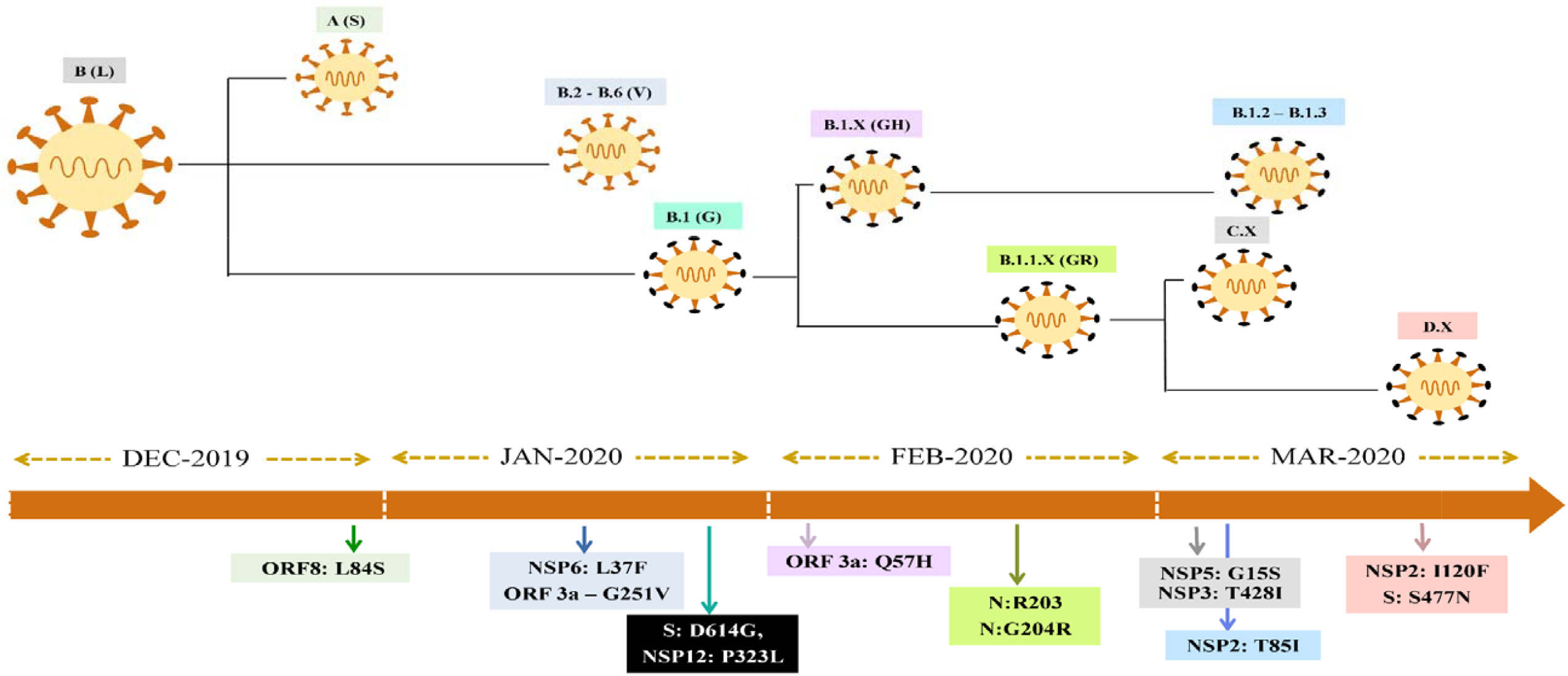
:Schematic representation of the major evolutionary events/ amino acid substitution that give rise to SARS-CoV-2 variants in sequential order

### Dominance of D614G variant

Two mutations have become consensus: D614G in S (nucleotide 23,403, A to G) and P323L (also known as P4715L) in nsp12 (nucleotide 14,143, C to T). These mutations were present in 80.5% of the sequences and have defined the B.1 lineage (G in GISAID classification). The widely discussed D614G variant is speculated to have been introduced in Europe at the end of January (EPI_ISl_422424) before becoming globally dominant. Genomes with D614G mutations were assigned as B.1 by PANGOLIN or GH/GR by GISAID. Notably, founder lineage B.1 and its sub lineages B.1.X, B.1.1.X, D.X and C.X that carry both D614G and P323L mutations have become the dominant variants across the world (87% of global collection as per CoV-GLUE as on 30th November 2020).

As the pandemic has progressed several other major substitutions affecting the protein structure have appeared. These are Q57H (nucleotide 25,563, G to T) in ORF3a, R203K + G204R combination (nucleotide 28,881, GGG to AAC) in Nucleocapsid and T85I (nucleotide 1059 C to T) in ORF1a. The region-specific sub lineages C.1, C.2, D.1 and D.2 were found to cumulatively harbour multiple mutations. Amino acid substitution such as T428I and G15S in ORF1a were reported in sub lineages C.1 and C.2, and S477N substitution in S protein along with I120F in nsp2, which specifically established the sub lineage D.2 (Figure 1).

### Structural analysis of SARS-CoV-2 mutants

The possible structural consequences of eleven lineage-defining missense mutations identified in this study were investigated. Among the mutations, three were considered as stabilizing the respective protein structure while six mutations were destabilizing (Table 1). The significance of these mutations in evolutionary selection cannot be solely predicted by ΔΔG, or change in free energy. Hence for a precise interpretation, correlation of ΔΔG, ΔΔS and N-H S^2^ (Supplementary Table S2) order parameter values of the proteins have been taken into account based on fine local-alterations in structures. All lineage-defining mutations except two have reduced the vibrational entropies of the proteins thereby decreasing the flexibility in the structures (Table 1).

**Table 1.** Lineage-defining SNPs and their impact on protein structures

| Protein | Lineage-defining Mutation | $\Delta\Delta S$ in kcal.mol<sup>-1</sup>.K<sup>-1</sup> | Change in dynamics | $\Delta\Delta G$ kcal.mol<sup>-1</sup>-DUET) | $\Delta\Delta G$ kcal.mol<sup>-1</sup>-SDM) | Stability |
| --- | --- | --- | --- | --- | --- | --- |
| Nsp12 | P323L | -0.33 | Decreasing flexibility | 0.43 (Stabilizing) | 1.57 (Stabilizing) | Stabilizing |
| Spike | D614G | -0.01 | Decreasing flexibility | 0.46 (Stabilizing) | 2.33 (Stabilizing) | Stabilizing |
| Orf3a | G251V | -0.39 | Decreasing flexibility | -0.6 Destabilizing | -2.19 Destabilizing | Destabilizing |
|  | Q57H | 0.44 | Increasing flexibility | -1.25 Destabilizing | 0.87 Stabilizing | Inconclusive |
| Orf8 | L84S | 0.30 | Increasing flexibility | -1.41 Destabilizing | -1.41 Destabilizing | Destabilizing |
| Nsp2 | T85I | 0.07 | Increasing flexibility | 0.54 (Stabilizing) | 1.93 (Stabilizing) | Stabilizing |
|  | I120F | -1.30 | Decreasing flexibility | -1.04 Destabilizing | -0.21 Destabilizing | Destabilizing |
| Nsp6 | L37F | -0.29 | Decreasing flexibility | -0.72 (Destabilizing) | -0.04 (neutral) | Inconclusive |
| Nucleocapsid protein (N) | R203K | -0.98 | Decreasing flexibility | -1.57 (Destabilizing) | -0.48 (Destabilizing) | Destabilizing |
|  | G204R | -0.16 | Decreasing flexibility | -1.06 (Destabilizing) | -1.95 (Destabilizing) | Destabilizing |
| Nsp5 | G15S | -0.31 | Decreasing flexibility | -0.98 (Destabilizing) | -0.79 (Destabilizing) | Destabilizing |

Additionally, the impact of mutations in key structural proteins that potentially allows any pathogen to escape available treatment and prevention regimes were investigated. Among the 59 major missense mutations, our analysis using both SDM and DUET server predicted 16 missense mutations as stabilizing 23 missense mutations as destabilizing the protein structure. Twenty major mutations were predicted to be neither stabilizing nor destabilizing as the ΔΔG values provided by SDM and DUET servers were contradictory (Table 2).

**Table 2:** Predicted effect of protein stability in the presence of amino acid mutations in the SARS-COV-2 genomes.

| Protein | Mutations | $\Delta\Delta G$ SDM<br>(kcal/mol) | $\Delta\Delta G$ DUET<br>(kcal/mol) | Stability |
| --- | --- | --- | --- | --- |
| Spike | A222V | 0.95 | 0.91 | Stabilizing |
|  | S477N | 0.31 | 0.02 | Stabilizing |
|  | L18F | -0.801 | -0.46 | Destabilizing |
|  | N439K | -0.29 | 0.3 | Inconclusive |
|  | L5F | -0.801 | -0.1 | Destabilizing |
|  | W1214G | -1.913 | -0.28 | Destabilizing |
|  | R21I | -0.856 | 0.46 | Inconclusive |
|  | A262S | -2.13 | -1.66 | Destabilizing |
|  | S98F | 1.23 | -0.58 | Inconclusive |
|  | D1163Y | 0.21 | 0.26 | Stabilizing |
|  | G1167V | -0.58 | -2.25 | Destabilizing |
|  | D936Y | -0.13 | -0.32 | Destabilizing |
|  | P272L | 2.12 | 0.36 | Stabilizing |
|  | D80Y | 0.77 | -3.08 | Inconclusive |
|  | E583D | -0.69 | -0.86 | Destabilizing |
|  | P1263L | -0.231 | 1.29 | Inconclusive |
|  | K1073N | -0.48 | -0.45 | Destabilizing |
|  | D253G | -0.19 | 0.04 | Inconclusive |
|  | T723I | 0.64 | 0.21 | Stabilizing |
|  | A688V | -0.18 | 0.03 | Inconclusive |
|  | A626S | -2.6 | -1.66 | Destabilizing |
|  | L54F | -0.61 | -1.33 | Destabilizing |
|  | H655Y | 0.6 | 1.44 | Stabilizing |
|  | G769V | 0.6 | 0.14 | Stabilizing |
|  | L176F | 0.12 | -0.95 | Inconclusive |
|  | G1124V | -1.52 | -0.14 | Destabilizing |
|  | V622F | -0.2 | -0.67 | Destabilizing |
|  | S255F | 0.94 | -0.8 | Inconclusive |
|  | H49Y | 0.4 | 1.14 | Stabilizing |
|  | D839Y | -0.389 | -1.08 | Destabilizing |
|  | V1176F | -0.92 | -0.55 | Destabilizing |
|  | D215H | 0.8 | 1.35 | Stabilizing |
|  | H146Y | 1.139 | -0.29 | Inconclusive |
|  | A879S | -2.57 | -1.69 | Destabilizing |
|  | Q677H | 0.98 | -0.48 | Inconclusive |
|  | D1084Y | -0.43 | -0.03 | Destabilizing |
|  | V1068F | -1.05 | -1.15 | Destabilizing |
|  | P25S | -0.392 | 0.93 | Inconclusive |
|  | A520S | -1.23 | -0.15 | Destabilizing |
|  | G261V | 0.16 | 0.16 | Stabilizing |
|  | D574Y | -0.56 | -0.45 | Destabilizing |
|  | T29I | 0.48 | 0.51 | Stabilizing |
|  | Y453F | -0.17 | -0.48 | Destabilizing |
|  | N501Y | 0.41 | -0.42 | Inconclusive |
|  | S939F | 0.76 | -0.71 | Inconclusive |
|  | T95I | 1.91 | 0.37 | Stabilizing |
|  | Q675H | 0.8 | -0.4 | Inconclusive |
| **Nucleocapsid** |  |  |  |  |
|  | A220V | -0.51 | -1.13 | Destabilizing |
|  | S194L | 1.15 | -0.02 | Inconclusive |
|  | D103Y | 1.45 | 0.55 | Stabilizing |
|  | P13L | 0.84 | 0.23 | Stabilizing |
|  | S197L | 1.27 | 0.26 | Stabilizing |
|  | A398V | -0.98 | -1.03 | Destabilizing |
|  | P199L | 1.27 | -0.21 | Inconclusive |
|  | M234I | 0.69 | 0.41 | Stabilizing |
|  | S188L | 1.21 | -0.06 | Inconclusive |
|  | S183Y | 0.05 | -0.71 | Inconclusive |
| **Membrane** | T175M | 0.69 | -0.26 | Inconclusive |
| **Envelope** | P71S | -0.03 | -2.35 | Destabilizing |

### Balance of stabilizing and destabilizing mutations

Overall, from both the datasets, 70 amino acid substitutions in SARS CoV-2 were tested for stability of which 19 were stabilizing, 29 were destabilizing and 22 showed inconclusive results. Computational prediction to understand the effect of amino acid substitutions in SARS CoV-2 revealed a balance of stabilization and destabilization of the proteins.

When checked for amino acid substitutions, the stabilizing mutation in S protein predicted an increase in the rigidity of its structure (Figure 3; Supplementary Figure S1). The increased rigidities of the structure may provide a stable conformation to the protein that may positively influence the binding of spike protein to ACE2 receptor (27). Major mutations D614G and S477N were located at potential epitope regions (Codons 469–882) with S477N particularly positioned in the receptor-binding domain (RBD) of the S protein (319 – 541).

**Figure 3.**
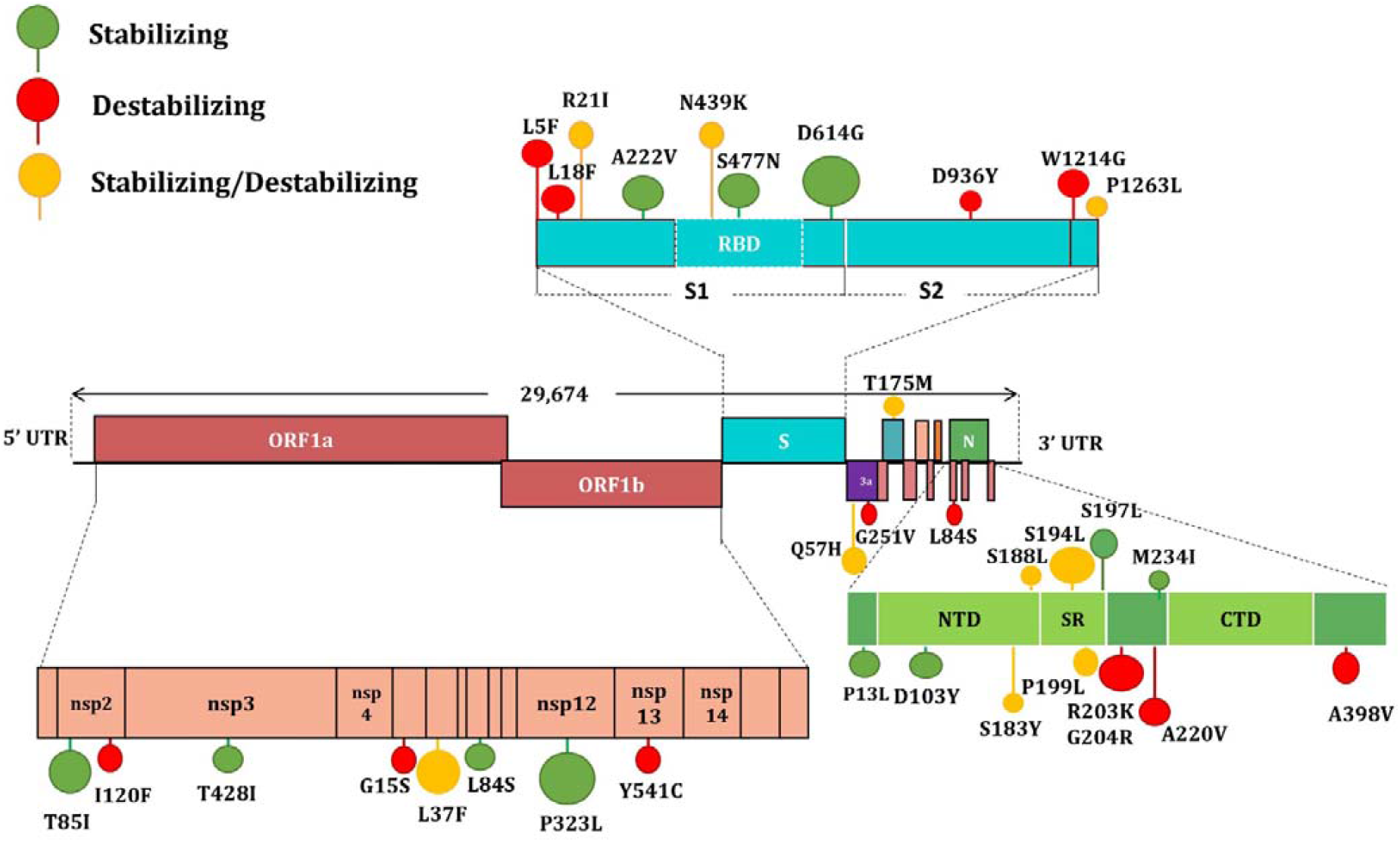
Schematic representation of SARS-CoV-2 genome organization, the major amino acid substitutions and stability of amino acid changes. Stabilizing mutations are coloured green, Destabilizing mutations are coloured red and mutations that neither stabilize nor destabilize are coloured in yellow.

Most frequent amino acid substitutions were observed in the N protein, in which the variants S194L, D103Y, P13L, S197L, M234I, and S188L were predicted to be stabilizing according to both the analytical servers (Table 2). In contrast, M and E proteins accounted for the least number of amino acid substitutions. The amino acid changes in M (T175M) indicated a stabilizing effect, while E does not account for any stabilizing variant. Structural analysis of double (D614G + S477N; D614G + A222V) and triple (D614G + S477N + A222V) mutation patterns in S protein indicated ΔΔG values of 0.228, 0.195 and 0.129, respectively (Table 3). This signifies that accumulation of spike mutation in D614G bearing lineages could potentially be affecting the stability of the spike and therefore may influence the binding affinity towards ACE2 receptor.

**Table 3:** Impact of double and triple mutation in the spike protein

| Protein | Combinations | Mutations | $\Delta\Delta G$ (pred) | C (pred) |
| --- | --- | --- | --- | --- |
| Spike | Independent | D614G | 0.422 | 0.892 |
|  | DOUBLE | D614G+S477N | 0.228 | 0.896 |
|  |  | D614G+A222V | 0.195 | 0.889 |
|  | TRIPLE | D614G+S477N+A222V | 0.129 | 0.129 |

## DISCUSSION

Since the beginning of COVID-19 pandemic, whole genome sequence based phylogenetic inference has been heavily utilized in tracing viral origins and transmission chains (28). However, as the virus has evolved with time, genomic data is being increasingly used in guiding infection risk and control strategies. Several genomic mutations have been mapped that seem to be of advantage to the virus (29). In parallel, numerous vaccine candidates have been designed using genomic data from the original SARS-CoV-2 strain of Wuhan and many are now approved for use or at late-stage trials (30, 31). Based on immunological data obtained from infected and recovered patients, rightly, almost all COVID-19 vaccine candidates of today are based on the original SARS-CoV-2 spike protein or its RBD domain (32 – 34). However, as vaccines are introduced and successful treatment options become available, it is vital that we carefully monitor the mutations in the immunogenic region of SARS-CoV-2 genome (35). Mapping these changes on protein structure will allow pre-emptive forecasting of the direction of change in vaccine effectiveness and guide future preparedness efforts. We analysed the impact of recurrent amino acid replacements in the genomic evolution and proteome stability of SARS-CoV-2 since its introduction in December 2019 to November 2020. Our analysis found an intriguing balance of stabilizing and destabilizing mutations, which may have allowed the SARS-CoV-2 to evolve and persist without losing pathogenicity.

SARS-CoV-2 is considered a slowly-evolving virus as it possesses an inherent proofreading mechanism to repair the mismatches during its replication. This is believed to have a crucial role in maintaining the stability and integrity of the viral genome (36, 37). Our analysis confirmed previously recorded positive natural selection of D614G, S477N (38), S477N, A222V and V1176F (39) variants and a global expansion of the PANGOLIN variant B.1 (11). In addition, we also observed a positive natural selection of Q57H (B.1.X), R203K/G204R (B.1.1.X), T85I (B.1.2-B.1.3), G15S+T428I (C.X) and I120F (D.X) variants (Figure 2).

Apart from the eleven clade defining mutations, some of the major missense mutations were in the four structural proteins (E, M, N and S). When analysed for their impact in the (*n=59*) respective protein structure, spike glycoprotein, more specifically its RBD domain, was found to be most vulnerable to frequent mutations. This may be due to the immunological observation that most neutralizing anti-SARS-CoV-2 antibodies have been found to target the RBD domain of the S protein (40,41). Consistent with this finding, a total of 4170 missense mutations have been reported in the spike protein, with 683 on the RBD domain alone (when CoV-Glue was accessed on 12th December 2020). Computational prediction to understand the effect of amino acid substitutions in E, M, N and S proteins revealed a balance of stabilization and destabilization of the proteins. While viral population carrying mutations with higher stabilizing effects (Positive ΔΔG values) would be expected to become the dominant variant, it is interesting to note that destabilization mutations in the major protein targets of SARS-CoV-2 have also generated variants that have been hugely successful. For example, many of the favourably selected variants such as L18F, L5F (Spike), R203K, G204R, A220V (Nucleocapsid) were found to be destabilizing the respective protein structure (Table 1). As destabilizing mutations are known for their crucial functional roles, a trade-off between stabilizing and destabilizing mutations may balance the protein function and structure in ways that are not fully understood yet (42,43).

In our study the effect of mutations on respective proteins was primarily estimated based on the physical change in free energy, on a single ‘native’ protein conformation. To allow the most robust correlation of mutations with the molecular evolution, the mutational effects when the protein is in unfolded state and possibility of structural adjustment of the folded state in response to the mutation needs to be explored in future studies as more structural dynamic information becomes available (44). While our study highlights the impact of ΔΔG analyses as a reference frame for evolutionary evaluation, molecular evolution is likely a consequence of complex amalgamation of changes in free energy, entropy, solvent accessibilities, etc (45). As the data on these unchecked parameters becomes available, predicting evolutionary selection of mutation with respect to the phylogeny would become confirmatory. Our study highlighting preliminary data linking free energy and phylogeny would help streamline the scope of future studies by providing a baseline matrix.

The currently circulating spike variants or RBD variants need to be taken into account while evaluating the vaccine candidate or neutralizing monoclonal antibodies against SARS-CoV-2 (46). Mapping the viral mutations that escape antibody binding is essential for accessing the efficacy of therapeutic and prophylactic anti-SARSCoV-2 agents (38, 47). Recently generated experimental evidence suggests that leading vaccines (mRNA-1273, BNT162b1 and ChAdOx1a) and two potent neutralizing antibodies (REGN10987 and REGN10933) are unlikely to be affected by the dominant variant D614G (32, 33, 48-50). As all three candidate vaccines encode RBD or the part of spike protein as antigens, the viral population is expected to try and escape by altering the positioning of the respective antigens (51) when vaccination induced selection pressure would be on. Notably, complete escape mutation map of 3,804 of the 3,819 possible RBD amino acid mutations against ten human monoclonal antibodies are already in place (38,51). The antigenic effect of key RBD mutations against REGN-COV2 cocktail (REGN10933 and REGN10987) showed N439K and K444R variants escaped neutralization only by REGN10987, while E406W escaped both individual REGN-COV2 antibodies and the cocktail (47). Similar strategies should be adopted to map all antibody resistance mutations against neutralizing antibodies elicited after vaccination. Once mutation escape maps are available for all successful vaccine candidates, vaccine roll out strategies should be carefully planned to counter geographically confined escape mutants.

## CONCLUSION

Our study highlights the importance of continued genomic surveillance, mutation mapping, stability analysis and potential escape mutation cataloguing in the pre- and post-vaccination period of SARS-CoV-2 in designing epidemiologically best vaccination programs. The currently observed mutation pattern and subsequent phylogenetic diversification of SARS-CoV-2 seem to be strongly influenced by the negative and positive selection pressures. The overall variation in SARS-CoV-2 sequences is currently low compared to many other RNA viruses. One of the possible reasons for the slow rate of mutations can be attributed to the widespread absence of neutralizing antibodies or the selective pressure. Once the virus population is challenged with the vaccine candidates or therapeutic monoclonal antibodies the currently known epitopes on surfaces of SARS-CoV-2 proteins are likely to undergo rapid forced change for survival. Thus, the prevalence of such possible escape mutations needs to be monitored even more carefully after vaccination if we are to remain ahead of this rapidly shifting pandemic curve.

## DATA AVAILABILITY

The genome sequences used in this is available in the Global Initiative on Sharing All Influenza Data (GISAID) with accession IDs (Supplementary Table S1)

## SUPPLEMENTARY DATA

Supplementary Data are available at online.

## ACKNOWLEDGEMENT

The authors would like to thank Mr. Soumya Basu (ICMR, Senior research Fellow) for his contribution and helpful advice in the structural analysis. The authors gratefully acknowledge the Department of Clinical Microbiology, Christian Medical College and Hospital, Vellore, Tamil Nadu, India, for providing all the necessary computational facilities for this work. We are grateful to the staff of Christian Medical College for their assistance with data curation.

## TABLE AND FIGURES LEGENDS

**Supplementary Figure S1:**
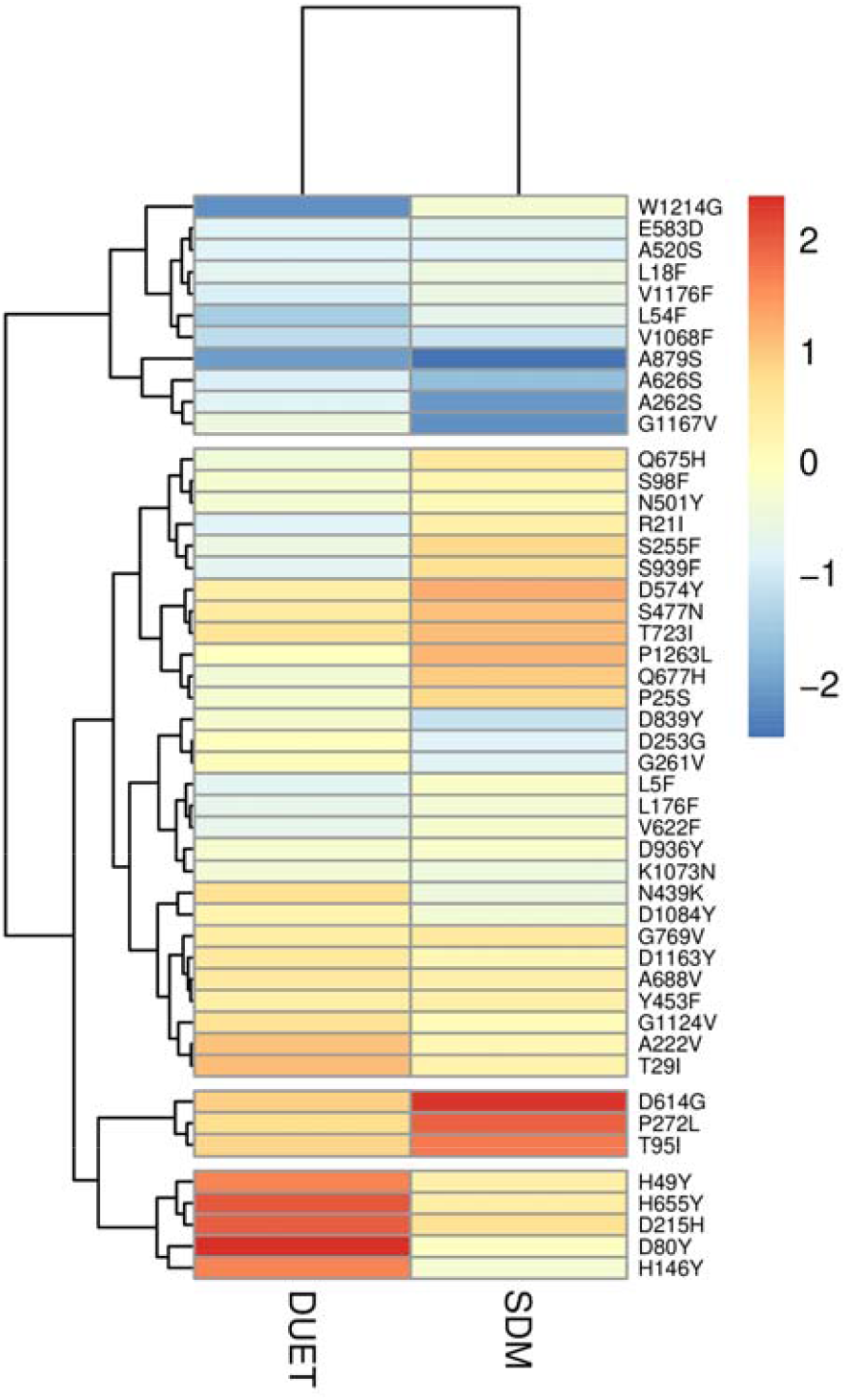
Heat map showing the stabilizing and destabilizing mutations of SARS-CoV-2 proteins based on the predicted ΔΔG values. The scale of heatmap ranged from −2 (Blue) to +2 (Red). Beige colour in the heat map indicates neutral ΔΔG values.

Supplementary Table S1: List of SARS-CoV-2 genome sequences downloaded from GISAID with accession IDs and metadata

Supplementary Table S2 : Residue level backbone stability values of lineage-defining mutations of SARS-CoV-2

## REFERENCES

1. World Health Organization. (2020) Coronavirus disease 2019 (COVID-19): situation report.

2. Worldometer, 2020 COVID-19 Coronavirus. Available at: https://www.worldometers.info/coronavirus/ (Cited date Dec 27, 2020).

3. World Health Organization: Rolling Updates on Coronavirus Disease (COVID19). (2020) Available at: https://www.who.int/emergencies/diseases/novel-coronavirus2019/events-as-they-happen. Accessed 18 May 2020.

4. Li, G. and De Clercq, E. (2020) Therapeutic options for the 2019 novel coronavirus (2019-nCoV). Nature., 149–150.

5. Hodgson, S. H., Mansatta, K., Mallett, G., Harris, V., Emary, K. R. and Pollard, A. J. (2020) What defines an efficacious COVID-19 vaccine? A review of the challenges assessing the clinical efficacy of vaccines against SARS-CoV-2. Lancet Infect Dis.

6. Sironi, M., Hasnain, S.E., Phan, T., Luciani, F., Shaw, M.A., Sallum, M.A., Mirhashemi, M.E., Morand, S. and González-Candelas, F. (2020) SARS-CoV-2 and COVID-19: A genetic, epidemiological, and evolutionary perspective. Infect Genet Evol., 84, 104384..

7. Singer, J., Gifford, R., Cotten, M. and Robertson, D. (2020) CoV-GLUE: a web application for tracking SARS-CoV-2 genomic variation. Preprints., 2020060225

8. Hu, B., Guo, H., Zhou, P. and Shi, Z. L. (2020) Characteristics of SARS-CoV-2 and COVID-19. Nature Rev Microbiol., 1–14

9. van Dorp, L., Acman, M., Richard, D., Shaw, L.P., Ford, C.E., Ormond, L., Owen, C.J., Pang, J., Tan, C.C., Boshier, F.A. and Ortiz, A.T. (2020) Emergence of genomic diversity and recurrent mutations in SARS-CoV-2. Infect Genet Evol., 83, 104351.

10. Dearlove, B., Lewitus, E., Bai, H., Li, Y., Reeves, D.B., Joyce, M.G., Scott, P.T., Amare, M.F., Vasan, S., Michael, N.L. and Modjarrad, K. (2020) A SARS-CoV-2 vaccine candidate would likely match all currently circulating variants. JPNAS., 117, 23652–23662.

11. Shu Y, McCauley J. GISAID: Global initiative on sharing all influenza data–from vision to reality. Eurosurveillance. 2017 Mar 30;22(13):30494.

12. Hadfield J, Megill C, Bell SM, Huddleston J, Potter B, Callender C, Sagulenko P, Bedford T, Neher RA. Nextstrain: real-time tracking of pathogen evolution. Bioinformatics. 2018 Dec 1;34(23):4121–3.

13. Rambaut, A., Holmes, E.C., Hill, V., OToole, A., McCrone, J., Ruis, C., du Plessis, L. and Pybus, O. (2020) A dynamic nomenclature proposal for SARS-CoV-2 to assist genomic epidemiology. Nat Microbiol., 5, 1403–1407

14. Tang, X., Wu, C., Li, X., Song, Y., Yao, X., Wu, X., Duan, Y., Zhang, H., Wang, Y., Qian, Z. and Cui, J. (2020) On the origin and continuing evolution of SARS-CoV-2. Natl Sci Rev., 7, 1012–1023

15. Plante, J.A., Liu, Y., Liu, J., Xia, H., Johnson, B.A., Lokugamage, K.G., Zhang, X., Muruato, A.E., Zou, J., Fontes-Garfias, C.R. and Mirchandani, D. (2020) Spike mutation D614G alters SARS-CoV-2 fitness. Nature.,1–6.

16. Korber, B., Fischer, W.M., Gnanakaran, S., Yoon, H., Theiler, J., Abfalterer, W., Hengartner, N., Giorgi, E.E., Bhattacharya, T., Foley, B. and Hastie, K.M. (2020) Tracking changes in SARS-CoV-2 Spike: evidence that D614G increases infectivity of the COVID-19 virus. Cell., 182, 812–827.

17. Hoffmann, M., Kleine-Weber, H., Schroeder, S., Krüger, N., Herrler, T., Erichsen, S., Schiergens, T.S., Herrler, G., Wu, N.H., Nitsche, A. and Müller, M.A. (2020) SARS-CoV-2 cell entry depends on ACE2 and TMPRSS2 and is blocked by a clinically proven protease inhibitor. Cell., 181, 271-280.e8

18. Katoh, K., Rozewicki, J., Yamada, K.D. (2019) MAFFT online service: multiple sequence alignment, interactive sequence choice and visualization. Brief Bioinform., 20, 1160–6.

19. Nguyen, L.T, Schmidt, H.A, Von Haeseler, A., Minh, B.Q. (2015) IQ-TREE: a fast and effective stochastic algorithm for estimating maximum-likelihood phylogenies. Mol Biol Evol., 32, 268–74.

20. Letunic, I. and Bork, P. (2019) Interactive Tree Of Life (iTOL) v4: recent updates and new developments. Nucleic Acids Res., 47, W256–W259.

21. Seemann T. Snippy: rapid haploid variant calling and core SNP phylogeny. GitHub. Available at: github.com/tseemann/snippy. 2015.

22. Pandurangan, A.P., Ochoa-Montaño, B., Ascher, D.B. and Blundell, T.L. (2017) SDM: a server for predicting effects of mutations on protein stability. Nucleic Acids Res., 45, W229–W235.

23. Pires, D.E., Ascher, D.B. and Blundell, T.L. (2014) DUET: a server for predicting effects of mutations on protein stability using an integrated computational approach. Nucleic Acids Res., 42, W314–W319.

24. Rodrigues, C.H., Pires, D.E. and Ascher, D.B. (2018) DynaMut: predicting the impact of mutations on protein conformation, flexibility and stability. Nucleic Acids Res., 46, W350–W355.

25. Goethe, M., Fita, I. and Rubi, J.M. (2015) Vibrational entropy of a protein: large differences between distinct conformations. J Chem Theory Comput., 11, 351–359.

26. Cilia, E., Pancsa, R., Tompa, P., Lenaerts, T., Vranken, W.F. (2014) The DynaMine webserver: Predicting protein dynamics from sequence. Nucleic Acids Res 42, 264–270.

27. Ou, J., Zhou, Z., Dai, R., Zhao, S., Wu, X., Zhang, J., Lan, W., Cui, L., Wu, J., Seto, D., Chodosh, J. (2021) V367F mutation in SARS-CoV-2 spike RBD emerging during the early transmission phase enhances viral infectivity through increased human ACE2 receptor binding affinity. BioRxiv.2020–03.

28. Oude Munnink, B.B., Nieuwenhuijse, D.F., Stein, M., O’Toole, A., Haverkate, M., Mollers, M., Kamga, S.K., Schapendonk, C., Pronk, M., Lexmond, P. and van der Linden, A. (2020) Rapid SARS-CoV-2 whole-genome sequencing and analysis for informed public health decision-making in the Netherlands. Nat Med., 26, 1802–1802.

29. Gómez-Carballa, A., Bello, X., Pardo-Seco, J., Martinón-Torres, F. and Salas, A. (2020) Mapping genome variation of SARS-CoV-2 worldwide highlights the impact of COVID-19 super-spreaders. Genome Res., 30, 1434–1448.

30. Dong, Y., Dai, T., Wei, Y., Zhang, L., Zheng, M. and Zhou, F. (2020) A systematic review of SARS-CoV-2 vaccine candidates. Signal Transduct Target Ther., 5, 1–14.

31. Alturki, S.O., Alturki, S.O., Connors, J., Cusimano, G., Kutzler, M.A., Izmirly, A.M. and Haddad, E.K. (2020) The 2020 pandemic: current SARS-CoV-2 vaccine development. Front Immunol., 11, 1880.

32. Corbett, K.S., Edwards, D., Leist, S.R., Abiona, O.M., Boyoglu-Barnum, S., Gillespie, R.A., Himansu, S., Schafer, A., Ziwawo, C.T., DiPiazza, A.T. and Dinnon, K.H. (2020) SARS-CoV-2 mRNA Vaccine Development Enabled by Prototype Pathogen Preparedness. bioRxiv.

33. Sahin, U., Muik, A., Derhovanessian, E., Vogler, I., Kranz, L.M., Vormehr, M., Baum, A., Pascal, K., Quandt, J., Maurus, D. and Brachtendorf, S. (2020) COVID-19 vaccine BNT162b1 elicits human antibody and TH 1 T cell responses. Nature., 586, 594–599.

34. Poland, G.A., Ovsyannikova, I.G. and Crooke, S.N. (2020) SARS-CoV-2 vaccine development: current status. Mayo Clin Proc., 95, 2172–2188

35. Li, Q., Wu, J., Nie, J., Zhang, L., Hao, H., Liu, S., Zhao, C., Zhang, Q., Liu, H., Nie, L. and Qin, H. (2020) The impact of mutations in SARS-CoV-2 spike on viral infectivity and antigenicity. Cell., 182, 1284–1294.

36. Robson, F., Khan, K.S., Le, T.K., Paris, C., Demirbag, S., Barfuss, P., Rocchi, P. and Ng, W.L. (2020) Coronavirus RNA proofreading: molecular basis and therapeutic targeting. Mol Cell., 79, 710–727

37. Bar-On, Y.M., Flamholz, A., Phillips, R. and Milo, R. (2020) SARS-CoV-2 (COVID-19) by the Numbers. eLife., 9, e57309

38. Starr, T.N., Greaney, A.J., Hilton, S.K., Ellis, D., Crawford, K.H., Dingens, A.S., Navarro, M.J., Bowen, J.E., Tortorici, M.A., Walls, A.C. and King, N.P. (2020) Deep mutational scanning of SARS-CoV-2 receptor binding domain reveals constraints on folding and ACE2 binding. Cell., 182,1295–1310.

39. Farkas, C., Mella, A. and Haigh, J.J. (2020) Large-scale population analysis of SARS-CoV2 whole genome sequences reveals host-mediated viral evolution with emergence of mutations in the viral Spike protein associated with elevated mortality rates. medRxiv.

40. Ju, B., Zhang, Q., Ge, J., Wang, R., Sun, J., Ge, X., Yu, J., Shan, S., Zhou, B., Song, S. and Tang, X. (2020) Human neutralizing antibodies elicited by SARS-CoV-2 infection. Nature., 584, 115–119.

41. Liu, L., Wang, P., Nair, M.S., Yu, J., Rapp, M., Wang, Q., Luo, Y., Chan, J.F.W., Sahi, V., Figueroa, A. and Guo, X.V. (2020) Potent neutralizing antibodies against multiple epitopes on SARS-CoV-2 spike. Nature., 584, 450–456.

42. Laha, S., Chakraborty, J., Das, S., Manna, S.K., Biswas, S. and Chatterjee, R. (2020) Characterizations of SARS-CoV-2 mutational profile, spike protein stability and viral transmission. Infect Genet Evol., 85, 104445.

43. Teng, S., Sobitian, A., Rhoades, R., Liu, D. and Tang, Q. (2020) Systemic Effects of Missense Mutations on SARS-CoV-2 Spike Glycoprotein Stability and Receptor Binding Affinity. Brief. Bioinformatics., bbaa233

44. Sikosek, T., Chan, H.S. (2014) Biophysics of protein evolution and evolutionary protein biophysics. J R Soc Interface. 11, 20140419.

45. Echave, J., Wilke, C.O. (2017) Biophysical models of protein evolution: understanding the patterns of evolutionary sequence divergence. Annu Rev Biophys. 46, 85–103.

46. Weisblum, Y., Schmidt, F., Zhang, F., DaSilva, J., Poston, D., Lorenzi, J.C., Muecksch, F., Rutkowska, M., Hoffmann, H.H., Michailidis, E. and Gaebler, C. (2020) Escape from neutralizing antibodies by SARS-CoV-2 spike protein variants. Elife., 9, e61312.

47. Starr, T.N., Greaney, A.J., Addetia, A., Hannon, W.H., Choudhary, M.C., Dingens, A.S., Li, J.Z. and Bloom, J.D. (2020) Prospective mapping of viral mutations that escape antibodies used to treat COVID-19. bioRxiv.

48. Voysey, M., Clemens, S.A.C., Madhi, S.A., Weckx, L.Y., Folegatti, P.M., Aley, P.K., Angus, B., Baillie, V.L., Barnabas, S.L., Bhorat, Q.E. and Bibi, S. (2020) Safety and efficacy of the ChAdOx1 nCoV-19 vaccine (AZD1222) against SARS-CoV-2: an interim analysis of four randomised controlled trials in Brazil, South Africa, and the UK. The Lancet.

49. Baum, A., Fulton, B.O., Wloga, E., Copin, R., Pascal, K.E., Russo, V., Giordano, S., Lanza, K., Negron, N., Ni, M. and Wei, Y. (2020) Antibody cocktail to SARS-CoV-2 spike protein prevents rapid mutational escape seen with individual antibodies. Science., 369, 1014–1018.

50. McAuley, A.J., Kuiper, M.J., Durr, P.A., Bruce, M.P., Barr, J., Todd, S., Au, G.G., Blasdell, K., Tachedjian, M., Lowther, S. and Marsh, G.A. (2020) Experimental and in silico evidence suggests vaccines are unlikely to be affected by D614G mutation in SARS-CoV-2 spike protein. npj Vaccines., 5, 1–5.

51. Greaney, A.J., Starr, T.N., Gilchuk, P., Zost, S.J., Binshtein, E., Loes, A.N., Hilton, S.K., Huddleston, J., Eguia, R., Crawford, K.H. and Dingens, A.S. (2020) Complete mapping of mutations to the SARS-CoV-2 spike receptor-binding domain that escape antibody recognition. Cell Host & ‘Microbe.

